# A pilot study of assisting IVF by personalized Endo-Gym^®^ exercises

**DOI:** 10.1101/2020.04.19.049379

**Authors:** Katalin Csehek, Peter Tompa

## Abstract

Assisted reproductive technologies (ARTs), especially *in vitro* fertilization (IVF) have revolutionized human reproduction technology, helping millions of subfertile couples to conceive and deliver a baby. IVF, however, is not an easy procedure, as treatment cycles incur heavy financial, physical and psychological burden, yet they result in live birth less than once in four attempts. Based on our experience with 251 women, many in their 40s, here we suggest that Endo-Gym^®^ method (for endocrine gymnastics), a combination of personalized physical exercises, fertility-optimizing diet and personal coaching, can significantly increase IVF success, probably by both reducing distress caused by repeated cycles and alleviating fertility-affecting problems, such as polycystic ovary syndrome (PCOS) and premature ovarian failure (POF). The program can also relieve other gynecological conditions, such as irregular or painful period, pelvic organ prolapse (POP) and incontinence, and is also often chosen by women as their regular fitness regime for general wellbeing. We provide detailed statistics of success in various conditions and suggest that distinct elements of Endo-Gym^®^ cooperate to exert positive physiological and psychological effects that help re-establish sexual hormone balance and boost reproductive fitness. We anticipate that further, controlled testing will enable to put the benefits of Endo-Gym^®^ on a rational basis and enable to introduce this approach as a beneficial complement of IVF, and maybe also other branches of ART.

## Introduction

Couples with fertility problems can resort to assisted reproductive technologies (ART) to boost their chances of conception and successful delivery. The best known and most generally applied ART is *in vitro* fertilization (IVF), in which, following hormonal stimulation, oocytes retrieved from the ovary of subfertile women are inseminated in a test tube and then transferred to the uterus [1, 2]. IVF was implemented in 1978, and since then millions of subfertile couples have used this and related ART technologies, resulting in an estimated number of 8 million babies worldwide [3]. It is anticipated that maybe 3% of world population may come from births assisted by IVF by year 2100. This tremendous success is also the result of continuous research that has brought regular updates and improvements in IVF technology, such as cryopreservation of surplus embryos, egg donation to women with empty ovaries, intracytoplasmic sperm injection to treat male factor infertility and uterine transplantation, to name just a few [1]. This rapid technological advance forecasts that the repertoire of possible IVF approaches will keep broadening, both on the social and scientific side, in the future.

It should not be neglected, however, that despite its sweeping success and broad applications, IVF is not a routine and comfortable procedure, as it puts a substantial financial, physical and psychological burden on couples. Notwithstanding all the advances in technology, almost 80% of patients who undergo an IVF cycle will not eventually have a baby, and often cycles have to be repeated multiple times for conception and live birth (2.7 cycles on the average, with about 38% overall success after 3 cycles) [1]. Due to the frustration of being unable to conceive naturally, the very decision of undertaking infertility treatment is already difficult [4], and repeated unsuccessful cycles put the couples through enormous stress and maybe long-term negative effects [5]. Evidence points to that the anxiety and distress experienced may negatively affect the outcome of IVF itself [6, 7] and result in the termination of the cycle before conclusion. About 20% of couples drop out after the first or second IVF cycle, with the primary reason being the psychological burden (and high cost in certain countries), which probably explains why “milder” treatment regimens usually entail less dropout [8, 9].

The initial stressful decision to undergo IVF, subsequent dropout and uncertainty in waiting periods are important concerns for age, as the average age of women undergoing IVF is 35.5 years, close to 40 where IVF success starts to decline sharply [1]. Therefore, although IVF – and other ART technologies – constantly evolve, improvements to reduce physical and emotional burden and to increase success rate are always welcome. Prior prognosis of success is also of paramount importance for counseling couples. The most widely accepted approach is ovarian reserve testing (ORT), which can not only provide an indicator of potential IVF success, but may also suggest individualized treatment protocols [10]. As currently there are limited options to improve poor ovarian reserve prior to IVF [11], exploring alternative methods may be highly beneficial.

With this goal in mind, here we report on a pilot study with Endo-Gym^®^ (endocrine gymnastics) as a complementary approach for boosting physiological and psychological fitness before treatment, and potentially increasing the success rate of IVF. Beneficial effects of the method combining personalized physical exercises, fertility-optimizing diet and personal coaching are statistically established by following 251 women who have exercised Endo-Gym^®^ for various periods of time during 2012-2019, with the goal of improving their success in IVF or seeking remedy for a range of other gynecological conditions. Several of them had already undergone unsuccessful IVF cycles or struggled with directly related fertility issues, such as polycystic ovary syndrome (PCOS) [12] or premature ovarian failure (POF, also ovarian insufficiency) [13]. Others had different but not less disturbing gynecological problems, such as incontinence, pelvic organ prolapse (POP), irregular/painful periods and low libido. Many women also opted for Endo-Gym^®^ simply to boost their fitness, i.e. general physical/mental wellbeing. We have assessed the experience of the women by analyzing questionnaires collected before, during and after the treatment. Our statistical analyses and individual case studies show that Endo-Gym^®^ can: (i) provide a transition period facilitating the initial decision of undergoing IVF, (ii) alleviate stress and physical burden during hormonal stimulation, (iii) help increase the chance of conception and live delivery, and (iv) provide complementary remedy for a broad range of other related or unrelated gynecological conditions.

In all, this pilot study provides the proof of concept that Endo-Gym^®^ helps couples undergo successful IVF and it can also relieve other problems. Whereas the mechanisms underlying its effects are not fully clear, we perceive that the exercises positively affect both mental wellbeing and reproductive fitness. Our long-term goal is to provide thorough condition-specific evidence in further, more extended, longitudinal studies.

## Materials and methods

### Institute for Endocrine Gymnastics

The research outlined in this study has been conducted for 8 years (2012-2019) based on the principles that culminated in the formulation of Endo-Gym^®^ method and the foundation of the Institute for Endocrine Gymnastics (IEG) in 2018 (https://www.instituteforendocrinegymnastics.org/). The underlying idea is that a complex and dedicated program based on cutting-edge research can offer integrated, personalized treatment options for women of reduced fertility and a variety of gynecological conditions, naturally relieving stress symptoms, re-establishing sexual hormonal balance and improving mental wellbeing, providing a natural and beneficial complement to IVF.

With trained professionals, the ambition of the Institute IEG is to: (i) raise funds to conduct research on improving reproductive fitness, (ii) serve as a center of integrated exercises, (iii) extend possible complementary treatments to other gynecological conditions, (iv) educate the public about the importance and relevance of hormonal functions in coping with daily challenges, and (v) advance awareness of the importance of bridging scientific understanding and natural, integrated approaches for troublesome gynecological problems.

### Endo-Gym^®^ method

The study is based on the Endo-Gym^®^ method [14], an integrated, complex system that combines regular, personalized physical exercises, fertility-optimizing diet [15-19] and personal coaching through an extended period. As the explicit ambition of the program is to regularize hormonal balance, we term it endocrine gymnastics, Endo-Gym^®^.

The exercises incorporate elements of High Intensity Interval Training (HIIT) which was found to improve metformin 72-h glucose control by reducing the frequency of hyperglycemic peaks [20]; Aviva Dance steps that is under validation in terms of its effect on increasing blood circulation in the female reproductive organs [21]; breath-focused yoga practices that are found to decrease cortisol levels [22]. Fertility-optimizing diet emphasizes limiting blood sugar spikes by altering eating habits when it comes to quality and volume of carbohydrates, also considering gluten elimination when required, regulating the intake of phytohormones, consuming sufficient fiber to improve estrogen metabolism and minimizing xenoestrogen exposure [23, 24]. Personal coaching included regular consultations and adaptation of the program motivated by personal needs, potential physical problems and individual advance of the client. Personalization has particular relevance in case of additional health problems, such as lower back, knee or shoulder injuries, and to respond to the changing needs according to the phases of menstrual cycle.

These diverse elements have been combined to develop a complex and personalized system, which constitutes elementary units (cycles) of 12 sessions over a period of 16 weeks. Each session combines half an hour personal consultation and half an hour supervised exercise. Sessions are usually complemented by shorter individual daily home practices, typically 2 × 4min a day. There is a broad distribution of the number of cycles completed by clients, most of them completing at least three (termed “regulars”), and many of them following the exercises for years.

### Applying Endo-Gym^®^

The research outlined in this paper has been conducted in Brussels for close to a decade. Women engaged in the program came through advertisements or personal contacts, and had most often already undergone some medical treatment or IVF with unsatisfactory result. They were always accepted in the program and their treatment commenced with a personal interview based on filling out a Health History Questionnaire (HHQ) assembled by the Institute for Integrative Nutrition and a Hormone Assessment Questionnaire (HAQ), as recommended by Sarah Gottfried MD [25]. Based on their health history, outcome of the questionnaires and consultations, a personalized program was compiled, including (i) targeted physical exercises [14], (ii) cycle-adapted personal diet [15-19] and (iii) individual coaching.

### Monitoring progress and analyzing data

HHQ and HAQ were also filled out by clients during the program and at the end, and progress was monitored by comparing answers in the three questionnaire. Success was measured by improvements in personal accounts, often supported by medical diagnoses (formal definitions in the different categories are given in Table 1). Occasionally (although not systematically) progress was also assessed by anecdotal observation/evidence of outstanding cases, a few of which are mentioned in Results.

**Table 1.** Definition of gynecological problems and their measure of success in Endo-Gym^®^ exercises. The table summarizes the various diagnosis or symptom(s) associated with gynecological problems that prompt Endo-Gym^®^ exercises. Success is measured as percent of women reporting a medically diagnosed or perceived improvement in their condition, as outlined in four different categories.

| **class** | **diagnosis/symptom** | **definition/measure of success** |
| --- | --- | --- |
| **Baby** |  |  |
|  | subfertility unexplained | successful pregnancy (and delivery)<sup>a</sup> |
|  | Polycystic ovary syndrome (PCOS)/premature ovarian failure (POF)/thyroid/endometriosis | successful pregnancy (and delivery)<sup>a</sup> |
| **Hormonal** |  |  |
|  | menstrual problem (irregular/painful) | regularization of menstrual cycle, mitigation of pain<sup>b</sup> |
|  | PCOS | regular menstruation and/or ovulation (confirmed by test strips)<sup>a,b</sup> |
|  | POF | observable oocytes in an empty ovary<sup>a</sup> |
|  | menopause | relief of symptoms such as dryness, hot flushes, insomnia<sup>b</sup> |
|  | low libido (low sexual desire) | Improved sexual desire (sometimes through feedback from husband)<sup>b</sup> |
| **Physical** |  |  |
|  | pelvic organ prolapse (POP) | relief of discomfort<sup>b</sup> |
|  | muscle tone/incontinence | no sanitary pads are needed any more<sup>b</sup> |
| **Fitness** | general wellbeing | positive changes in fitness and mode<sup>b</sup> |

<sup>a</sup>success based on medical diagnosis
<sup>b</sup>success based on self-reporting

Formally, replies in HHQ and HAQ were entered in an anonymous database in which women were entered by their entry number into the program (1-251) and age, to comply with GDPR on handling personal data. We also recorded individual observations. In case of multiple observations, statistics was calculated and, when appropriate, compared to age-matched groups in large-scale IVF reports [1, 2, 26]. Striking anecdotal observations were also recorded and indicated above the background of given reference groups.

## Results

### Women engage in Endo-Gym^®^ with different gynecological problems

Women joining the Endo-Gym^®^ program often had a medical (gynecological) condition, for which they had been treated with unsatisfactory results. Many of them had already undergone unsuccessful IVF cycles and either were advised to drop out or simply wanted to improve their chances in future IVF attempts. To help advance and adapt treatments, they were put into four general target categories (Table 1). In category 1 (“Baby”), women aimed to conceive and deliver a baby either by natural intercourse or by returning to IVF. Their condition causing subfertility and compromising conception was often uncharacterized. In category 2 (“Hormonal”), women had conditions related to menopause or sub-fertility, such as POF (also empty ovary or ovarian insufficiency), early menopause, irregular/painful period or PCOS [12, 13] and low libido. Their primary aim was to relieve adverse long-term consequences of their condition (e.g. unwanted hair-growth, diabetic complications and potential cancer in PCOS and osteoporosis, joint problems, blood clotting and vaginal dryness in early menopause), rather than conceiving a baby. The third category (“Physical”) came with different but not least disturbing gynecological problems originating more in compromised muscle tone, causing incontinence and/or pelvic organ prolapse (POP). Finally, in the fourth category (“Other”), women sought regular exercises to boost their physical fitness and mental wellbeing. Our general perception is that besides particular goals with their given condition, practically all women wanted to increase their comfort and confidence of proper sexual functioning in their relationship. For each category, we defined different measures of success, which was monitored based on either medical diagnosis or self-reporting (Table 1).

### Success can be achieved at all ages with Endo-Gym^®^

Some of our clients already tried “alternative” approaches before they enrolled in an Endo-Gym^®^ group. Due to this prior history and frustrating attempts, their age distribution shows a bias to advanced age, as compared to IVF clients (Figure 1, Table 2). The average age of our clients when they started the program was 40.9y, with 53.0% (133/251) of them being 40y or older. As women tend to attempt IVF at an ever increasing age (Figure 1A), in a sense we have encountered a representative sample of future IVF clients. Women in the Baby category have a mean age of 38.2y, whereas in all other categories combined, 42.2y. In these latter groups, age distribution is heterogeneous, reflecting the distinct age of women with different problems (Figure 1, Table 2). Our experience with these women shows significant success (for definitions, cf. Table 1), on the average, 30% (Baby), 33.6% (Hormonal), 35.2% (Physical) and 82.5% (Other). For sub-categories in the four major categories, success rate varies and can reach very high values (e.g. 72.7% for PCOS in Hormonal group, cf. Table 2). As another measure of success and inspiration of the program, we may cite the strong tendency of women to stay on Endo-Gym^®^ beyond the first cycle. Initially, women always join the program for one cycle (12 sessions), but in a high proportion of the cases (between 55% and 65%, cf. Figure 1B) they remain for at least three cycles (became regular) and often take it on as a lifestyle change, staying on the program for years.

**Table 2.** Success of Endo-Gym^®^ in various gynecological conditions. The table enlists the type of problems with which women attend Endo-Gym^®^ exercises (statistics for 2012-2019). Women in “Baby” category sought to fall pregnant. The reason of their subfertility is most often unexplained. Women also come with many other, indicated gynecological conditions, which we placed into two not clearly separable categories “Hormonal” or “Physical”. We have also had many clients who chose Endo-Gym^®^ exercises as their regular physical activity for general wellbeing (“Fitness”). We indicate the number of women who attended the course in the different categories and indications, their mean ages and if they have decided to stay permanently on the program (regular practice: at least three cycles). The success rate of their program is calculated based on feedback (see Materials and methods), as defined in the different categories in Table 1 (for example, in Baby category, we define primary success as pregnancy).

| class | diagnosis/symptom | number | mean age | regular practice | success % |
| --- | --- | --- | --- | --- | --- |
| **Baby** |  |  |  |  |  |
|  | subfertility unexplained | 74 | 38.2 | 46 | 29.7 |
|  | PCOS/premature ovarian failure (POF)/ thyroid/endometriosis | 10 | 38.2 | 8 | 30 |
| **Hormonal** |  |  |  |  |  |
|  | menstrual problem (irregular/painful) | 54 | 38.8 | 30 | 27.7 |
|  | PCOS | 11 | 34.8 | 11 | 72.7 |
|  | POF | 12 | 42.6 | 10 | 66.7 |
|  | menopause | 31 | 50.7 | 10 | 32.2 |
|  | low libido (low sexual desire) | 41 | 41.0 | 26 | 19.5 |
| **Physical** |  |  |  |  |  |
|  | pelvic organ prolapse (POP) | 3 | 45.5 | 3 | 100 |
|  | muscle tone/incontinence | 85 | 42.5 | 45 | 32.9 |
| **Fitness** | general wellbeing | 62 | 42.6 | 40 | 82.2 |
| **Total** |  | **251** | **40.9** | **134** |  |

**Figure 1.**
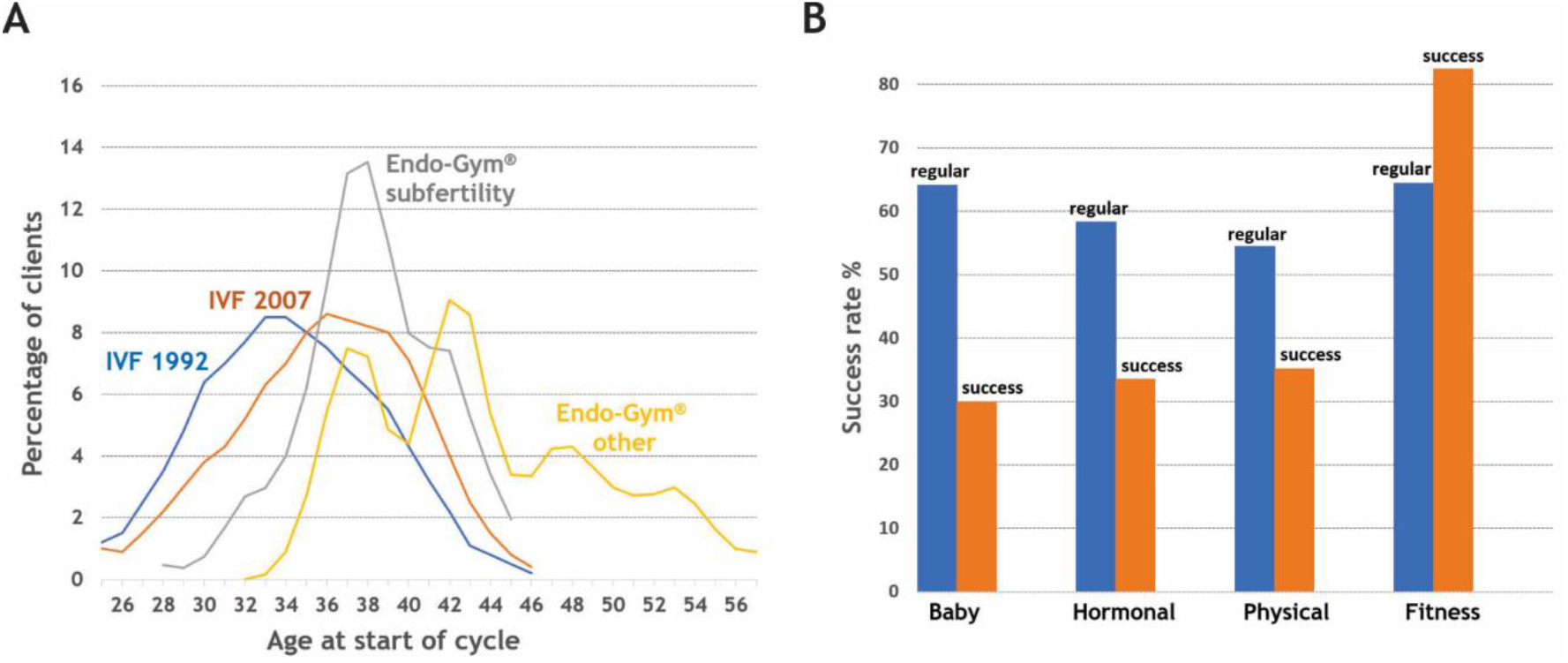
Basic statistics of attending IVF and Endo-Gym^®^. **(A)**Age distribution of women entering IVF treatment tends to drift to older age, as shown by comparing statistics for 1992 and 2007 (with mean 33.8y in 1992 vs. 35.5y in 2007, the same as in 2017 [1]). The age distribution of women attending Endo-Gym^®^ for conceiving a baby (subfertile) is skewed to even higher age (mean 38.2y). Women exercising Endo-Gym^®^ for other reasons (cf. Table 1) tend to be even older (mean 42.2y) and show multivariate distribution, apparently corresponding to the diverse underlying gynecological conditions. **(B)**A typical Endo-Gym^®^ cycle lasts for 12 sessions. Many women stay much longer on the program (complete at least 3 cycles, i.e. become “regular”). In all different categories, the exercises bring significant success (cf. Table1 for the definition of “success” in the various categories and Table 2 for detailed statistics).

### Complementing IVF in conception and delivery

As our primary ambition with Endo-Gym^®^ has been to help subfertile women to conceive and deliver a healthy baby, we paid particular attention to assess the success of this group (Baby, cf. Table 2 and Figure 2). We consider these values in light of statistics of IVF birth rate per treatment cycle (PTC), which is about 28% for women below 40y and is only about 4% for women above 40 (21% overall) [1] (cf. Figure 2A). In our case, with a total of 84 women undertaking Endo-Gym^®^ for pregnancy (Table 2), we observed a notable rates of success (Figure 2B). Of these women, 31.2% successfully conceived and 29.8% delivered a baby, which increased even more if they attended the exercises for at least three cycles (regulars, cf. Figure 2B). Another comparison also attests to the strength of Endo-Gym^®^: as suggested, women undertake these exercises at an older average age, yet their success of falling pregnant (34.5% vs. 26.9%) and delivering a baby (32.8% vs. 23.1%) does not drop drastically at the 40y border. Probably arguing for the strong synergy of Endo-Gym^®^ and IVF is the fact that many of our clients stayed on, or returned to, IVF while on Endo-Gym^®^ program, thus often their success can be attributed to the complementary effect of the two treatments.

**Figure 2.**
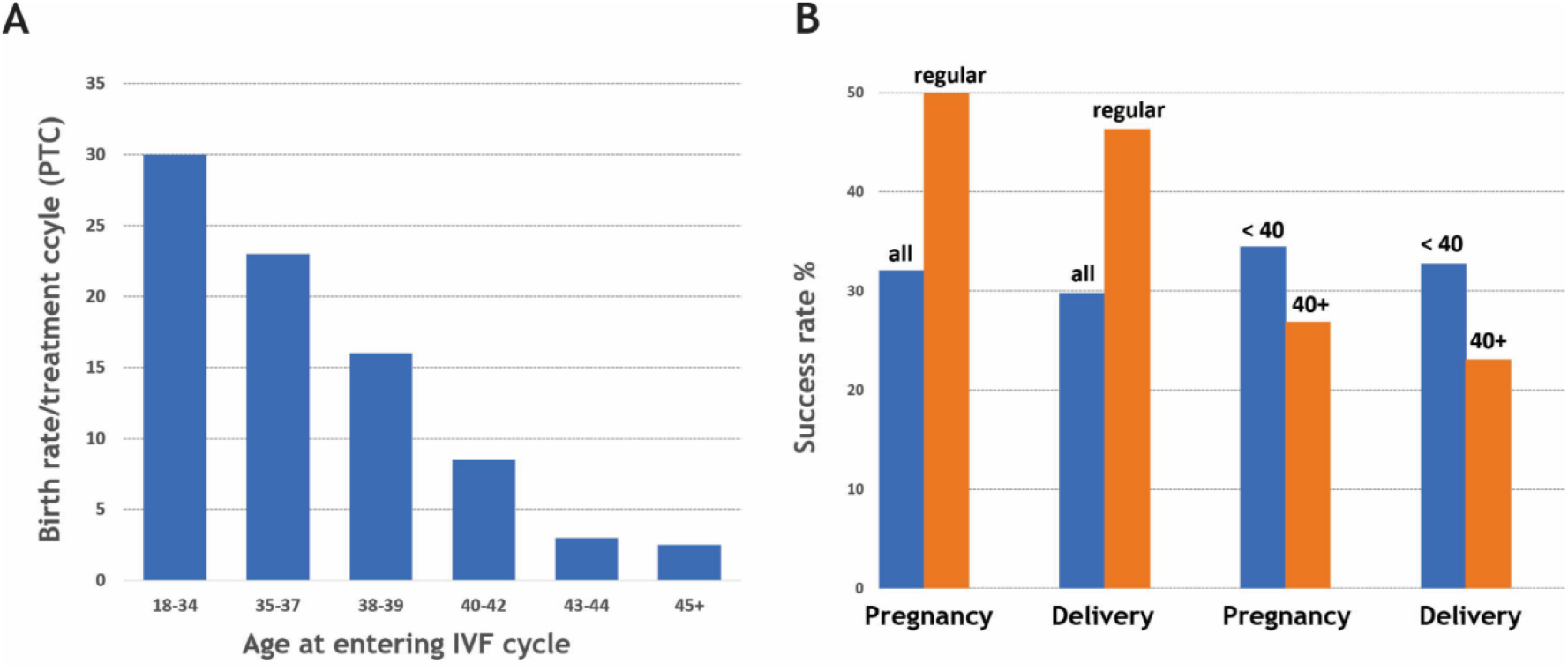
Success rates of IVF and different categories in Endo-Gym^®^. **(A)**The success of IVF (% live birth per treatment cycle, PTC) is around 21% overall, declining strongly with age, dropping to 4% above 40y of age (statistical data taken from ref. [1]). **(B)**Women who wish to conceive and deliver (Baby) show significant success upon undertaking Endo-Gym^®^ exercises (either through or outside an IVF program). Regular practice (at least 3 cycles, as opposed to “all” clients) entails a significantly elevated success rate, both in terms of pregnancy and successful delivery of a baby. Whereas age is a strong negative factor, women show significant success in both pregnancy and delivery (shown for all) even at or above 40y of age (40+).

### Notable anecdotal cases highlight efficacy in diverse conditions

On top of these statistical comparisons, we have also encountered several remarkable individual cases, which cannot be substantiated by statistics, yet they are very inspiring for clients considering to attend the program and IEG professionals aiming to promote Endo-Gym^®^. In accord, our plan is to corroborate these findings in large-scale studies in the future.

*Case 1:* one of the most remarkable cases is a woman of 39 years, who came to the exercises following a diagnosis of empty ovaries, having been proposed to undergo IVF with an egg donor. Remarkably, even within the first cycle of exercises (actually, after two months), she was found to have antral follicles growing, and could return to normal IVF with her own oocytes.

*Case 2:* a somewhat older woman started missing her period at 46, and was checked by a gynecologist for her follicle count and hormone levels. Because she was diagnosed with empty ovaries, further based on her hormone levels she was concluded to having entered menopause. She started to exercise, and in 6 months of Endo-Gym^®^, her regular periods returned.

*Case 3:* one client missed her cycles for two years, between 40 and 42. During this period, she sometimes received Clomid treatment, during which she occasionally observed weak bleeding (spotting). She then enrolled into the Endo-Gym^®^ program, and in four months’ time she produced her first proper menstruation again, which repeated in the next 2 months (she is still on the program).

*Case 4:* a woman of 40y had the opportunity to follow up her progress with regular blood tests. In February 2018, her follicle-stimulating hormone (FSH) was 249 IU/L, accompanied by missing periods. Her gynaecologist found that she also had POF. We started working with her in June, in a month’s time her FHS went down to 117 IU/L, followed by 52 IU/L in two months and by December, when she had already her second consecutive menstruation, her FSH was at 31.4 IU/L; furthermore her gynaecologist saw antral follicles growing again in her ovaries.

*Case 5:* as an additional touching story, we had a client of 40y who came off the contraceptive pill after 21 years. Her cycle stopped, and in three months she went to a homeopathic doctor. She received an oral homeopathic medication, but in half a year she could produce only two menstruations. She then started Endo-Gym^®^ exercises on a daily basis, and her normal cycle returned after a month. She eventually conceived and delivered a baby by IVF (due to the very low sperm count of her 61 year old husband).

*Case 6:* as a final notable case, we note a very young girl of 19y, who was rather desperate as she had not yet had a menstruation. She started the program, and in three weeks she had her first period ever. She felt a bit tired during the first days, then her energy level went back and she restarted exercising again. In three weeks she had another period, her follow up is still in progress.

## Discussion

The focus of this paper is to collate and analyze data on our experience with women with fertility problems and other lifestyle-related, disturbing gynecological indications. As outlined, our experience with a large number of women of rather advanced age (mean: 40.9y) is very positive, and anecdotal evidence is also suggestive of the favorable outcome of the treatment. The high percentage of women who stay on the program and practice Endo-Gym^®^ as their routine and regular fitness and wellbeing exercise confirms its general mental and physical benefits. In all, also considering that many of the clients only stop exercising Endo-Gym^®^ with the Institute because they change home or drastically change lifestyle, job, etc., all these are also suggestive of the positive personal experience it brings.

Its positive effect in a broad range of gynecological conditions speaks for itself. The high success rate of conceiving and delivering a baby gives hope for subfertile couples, whether without or with concomitant/subsequent IVF treatment. As its success rate, especially above 40y of age, is above that reported in IVF (Figure 2), it is not farfetched to consider and suggest Endo-Gym^®^ as a useful complement to IVF. Actually, women reported that if they started the exercises for assisting IVF, it helped them not only physically but also emotionally, to endure the stress concomitant to repeated cycles. As stress/psychological burden is the primary cause of dropout from the IVF program [8, 9], this may be another reason to suggest that Endo-Gym^®^ may help women start IVF and/or stay on the IVF program. As couples reported that due to the frustration of their inability to conceive naturally, even deciding on IVF is not easy [4], initial Endo-Gym^®^ exercises may help bridge this difficult time of decision, which may than be further aggravated by later positive effects.

In terms of its mechanism of action, Endo-Gym^®^ undoubtedly has strong, positive physiological effects. The exercises are made up of dozens of dynamic, stretching and relaxing moves, using a variety of breathing techniques, all designed to move not only core muscles, the mesentery and reproductive organs but also to pump blood to the pelvic cavity and its organs. We are all aware that regular exercises strengthen skeletal muscles [27] by incumbent hyperemia, i.e. an increase in skeletal muscle blood flow. Vasodilation in muscle arteries, however, is also accompanied vasoconstriction (reduced blood flow) in peripheral organs (kidneys, intestines, non-exercising skeletal muscles), to direct blood flow to contracting skeletal muscles. As our exercises are targeted at core muscles and reproductive organs, they probably reverse the underlying regulatory mechanisms of the sympathetic nervous system, causing vasodilation in core muscles and reproductive organs, increasing their blood flow, metabolism, neuronal activity and hormonal responsiveness.

Another strong component may come from psychosocial elements. Self-determination theory (SDT) contends that humans function effectively if they can satisfy three psychological needs: autonomy, competence and relatedness [28]. *Autonomy* derives from the perception that we are the origin of our choice, *competence* builds on our mastery – understanding and control – of our action, whereas *relatedness* pertains to being attached and respected by others. It can be hardly argued that competence and relatedness are missing in a routine IVF treatment, which are, however, inherent to the exercises of Endo-Gym^®^. Personal coaching provides a sense of understanding what we are doing, whereas sharing the experience with women of similar fate disperses the sense of isolation and provides continuous self-reinforcement, with immense and apparent added values. Last but not least, the diet suiting the purpose of resetting hormonal balance dysregulated around peri-menopause, menopause and under the conditions indicated (Table 1 and Table 2) also has great benefits.

As a final, interesting twist on our case with the benefits of the combination of these various components into this integrated system, it is notable that sometimes men (husbands) also attended our classes. Whereas their problems are definitely different, sometimes they also reported positive effects on erectile function (EF) and libido, and their presence and motivation had a very strong supportive and confirmatory effect on exercising women. This will definitely be a subject of future studies.

In conclusion, we are reporting on many positive effects of Endo-Gym^®^, which suggests that this integrated program may serve as a useful complement of IVF, and also as a valuable lifestyle component helping women improve their self-esteem by way of alleviating diverse gynecological problems and boosting their reproductive fitness and mental wellbeing. Of course, we are fully aware that our conclusions rely on a backward pilot analysis, which need to be – and will be – much strengthened by designed cohort studies. Our primary goal with the current report is to lay the foundation of such research in the near future.

